# *In vivo* assembly and trafficking of olfactory Ionotropic Receptors

**DOI:** 10.1101/441782

**Authors:** Liliane Abuin, Lucia L. Prieto-Godino, Haiyun Pan, Craig Gutierrez, Lan Huang, Rongsheng Jin, Richard Benton

**Affiliations:** Center for Integrative Genomics Génopode Building Faculty of Biology and Medicine University of Lausanne CH-1015 Lausanne Switzerland; Department of Physiology and Biophysics University of California, Irvine California 92697 USA

**Keywords:** ionotropic glutamate receptor, Drosophila, glycosylation, membrane trafficking, olfaction

## Abstract

lonotropic Receptors (IRs) are a large, divergent subfamily of ionotropic glutamate receptors(iGluRs), with roles in chemosensation, thermosensation and hygrosensation. Analogous to the synaptic targeting mechanisms of their iGluR ancestors, IRs are thought to form complexes of broadly-expressed co-receptors and selectively-expressed ‘tuning’ receptors to localise to sensory cilia. While tuning receptors’ extracellular ligand-binding domain (LBD) defines sensory specificity, the role of this domain in co-receptors is unclear. We identify a coreceptor-specific sequence in the LBD, which contains a single N-glycosylation site. Combining molecular genetic and cell biological analyses, we show that this site is dispensable for assembly of IR complexes in olfactory sensory neurons, but essential for endoplasmic reticulum exit of some,but not all, IR complexes. Our data reveal an important role for the IR co-receptor LBD in control of intracellular transport, provide novel insights into the stoichiometry and assembly of IR complexes, and uncover an unexpected heterogeneity in the trafficking regulation of this sensory receptor family.

## lntroduction

Ionotropic Receptors (IRs) are a subfamily of ionotropic glutamate receptors (iGluRs) [1], an ancient class of ligand-gated ion channels present in animals, plants and prokaryotes [2–4]. Comparative genomics analyses suggest that IRs evolved in the last common ancestor of Protostomes, and probably derived from the AMPA/Kainate clade of iGluRs [5], which have well-characterised roles in synaptic transmission in animal nervous systems [3]. In contrast to these iGluRs, IR repertoires have greatly expanded in size and display high sequence diversity within and between species [5]. Moreover, transcriptomic and *in situ* expression analyses in a range of animals indicate that *Ir* genes are expressed in peripheral, rather than central, neurons [5–9]. Functional analyses of IRs, in particular in *Drosophila melanogaster,* have shown that these receptors have diverse roles in environmental sensing, including in olfaction [6, 10–12], gustation [13–21], hygrosensation [22–24] and thermosensation [25, 26].

Within IR repertoires, two closely-related members, IR8a and IR25a, display several distinctive properties: first, they have the highest sequence identity and closest structural organisation to iGluRs [5], comprising an amino-terminal domain (ATD), a ligand-binding domain (LBD), and a transmembrane ion channel domain (Figure 1A). By contrast, most other IRs lack the ATD and have very low sequence identity to iGluRs, especially within the LBD (Figure 1A) [1, 5]. Second, these receptors are highly conserved, with unambiguous orthologues present across arthropods (for IR8a) or protostomes (for IR25a) [5]. Third, while many IRs are restricted to small populations of sensory neurons, IR8a and IR25a are broadly expressed across multiple, functionally-distinct neuron classes [1, 27]. Finally, genetic analysis in *D. melanogaster* indicates that loss of either IR8a or IR25a abolishes the responses of diverse sensory neuron classes [16, 19–23, 25, 27, 28]. These observations have led to a model in which IR8a and IR25a function as co-receptors that form heteromeric complexes with distinct sets of selectively-expressed, ‘tuning’ IRs, which determine the sensory response specificity of the complex [27].

**Figure 1.**
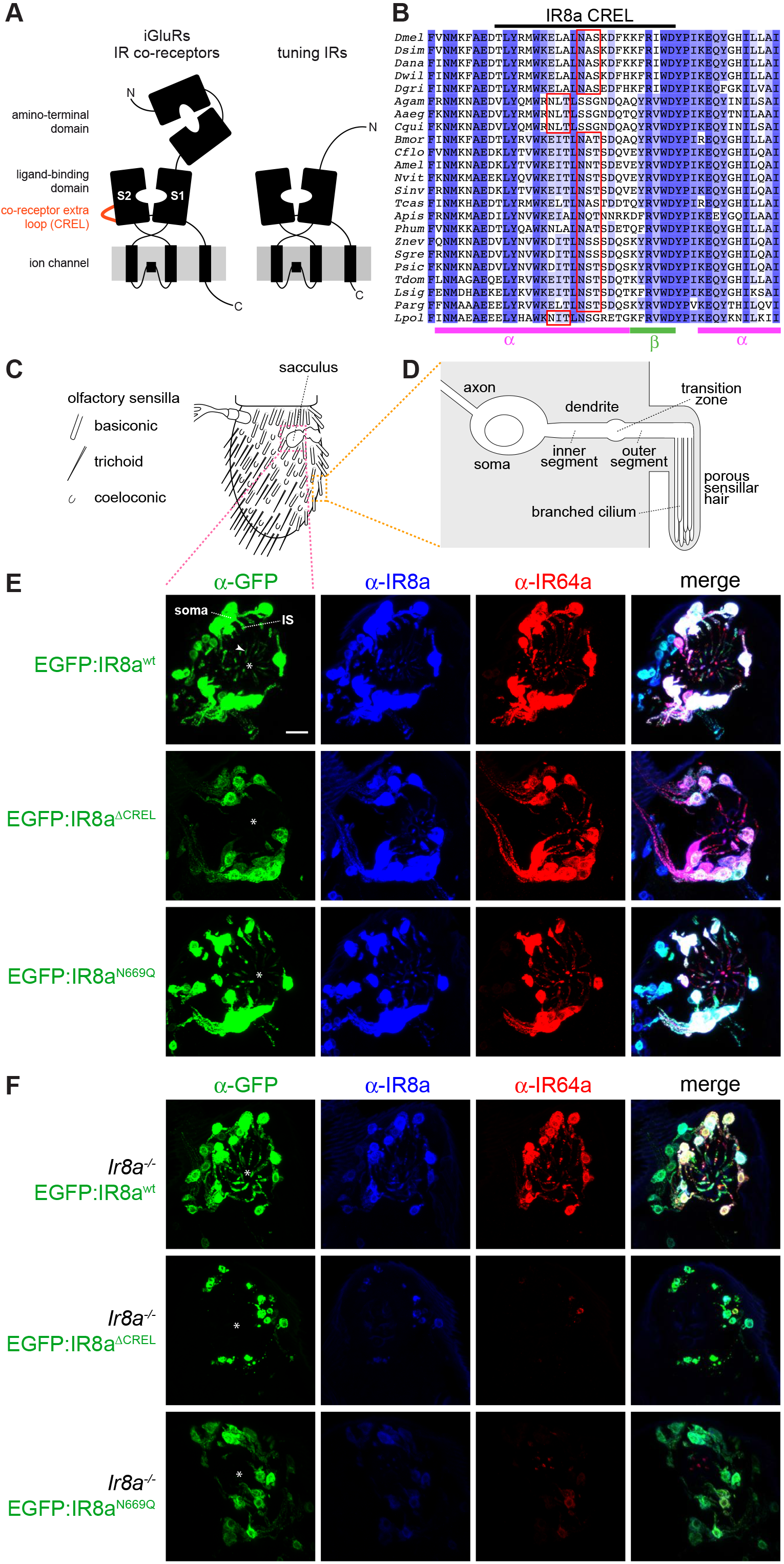
The IR co-receptor CREL functions in subcellular trafficking. (A) Schematic of the domain organisation of iGluRs, IR co-receptors and tuning IRs. (B) Alignment of the protein sequence spanning the CREL (co-receptor extra loop; black bar) in IR8a orthologues from the indicated species. Predicted N-glycosylation sites are highlighted with red boxes and predicted secondary structure is shown below the alignment. Species (top-to-bottom): *Drosophila melanogaster*, *Drosophila simulans*, *Drosophila ananassae*, *Drosophila willistoni*, *Drosophila grimshawi*, *Culex quinquefasciatus*, *Aedes aegpti*, *Anopheles gambiae*, *Bombyx mori*, *Tribolium castaneum*, *Camponotus floridanus*, *Solenopsis invicta*, *Apis mellifera*, *Nasonia vitripennis*, *Acyrthosiphon pisum*, *Pediculus humanus*, *Zootermopsis nevadensis*, *Schistocerca gregaria*, *Thermobia domestica*, *Phyllium siccifolium*, *Lepismachilis y-signata*, *Panulirus argus*, *Limulus polyphemus*. (C) Schematic of the *Drosophila* third antennal segment showing the distribution of different olfactory sensilla and other sensory structures. The orange box highlights a single sensillum shown in greater detail in (D). The pink box indicates the approximate field-of-view shown in panels (E-F), encompassing IR8a neurons in the sacculus that express the tuning receptor IR64a [12] (sensilla in the sacculus — which have a similar global organisation to surface sensilla as in (D) — are not schematised). (D) Schematic illustrating the main anatomical features of an olfactory sensory neuron (OSN); the morphology of the cuticular hair and the branched nature of the cilia vary between different sensilla classes (note: most sensilla contain more than one neuron per hair) [36]. (E) Immunofluorescence with antibodies against EGFP (green), IR8a (blue) and IR64a (red) on antennal sections of animals expressing the indicated transgenes in Ir8a neurons. Genotypes are of the form: *Ir8a-Gal4/UAS-EGFP:Ir8a*^*x*^. The white asterisks in (E-F) indicate the central cavity of sacculus chamber 3, into which the IR64a+IR8a-expressing OSN ciliated dendrites project. In the top left panel, the arrowhead marks the ciliated ending of one neuron; the soma and inner segment (IS) of this neuron are also indicated (the outer segment – before the cilium – is difficult to see because only trace levels of receptors are detected in this region). Scale bar (for all panels): 10 μm. (F) Immunofluorescence with antibodies against EGFP (green), IR8a (blue) and IR64a (red) on antennal sections of animals expressing the indicated transgenes in Ir8a neurons in an *Ir8a* mutant background. Genotypes are of the form: *Ir8a ^1^/Y;Ir8a-Gal4/UAS-EGFP:Ir8a^x^.* Both EGFP:IR8a^ΔCREL^ and EGFP:IR8a^N669Q^ fail to localise to the cilia in the absence of endogenous IR8a (the occasional projections from the soma represent protein within the inner segment only). Both proteins appear to be destabilised; consequently, endogenous IR64a is also detected at substantially lower levels in these two genotypes. OSNs that express EGFP:IR8a^ΔCREL^ also display signs of sickness (e.g., smaller soma).

While some progress has been made in defining the molecular basis of tuning IR response specificity [10, 11], the assembly and trafficking of IR complexes *in vivo* remain poorly understood. One intriguing unresolved question is the role of the LBD of the co-receptors. In heteromeric iGluR complexes, each subunit is thought to bind an extracellular ligand (typically glutamate or glycine) and to contribute to the gating of the ion channel pore [2, 3, 29]. It is unlikely that the IR co-receptors bind to the diversity of ligands that activate neurons in which they are expressed (which are presumed to be recognised by the LBD of the particular tuning IR in a given neuron type [10, 11, 27]). One possibility is that the IR co-receptor LBD interacts with a ligand that is present ubiquitously in the extracellular lymph fluid that bathes the ciliated outer dendritic segment of IR-expressing sensory neurons. The LBDs of both IR8a and IR25a retain most of the principal glutamate-contacting residues of iGluRs [1], raising the possibility that glutamate or a structurally-related molecule is such a ligand. However, the observation that several of these residues are dispensable for the function of IR8a [27] implies that the co-receptor LBD has a role that is independent of ligand interactions.

## Results and Discussion

### lR co-receptor LBDs contain a unique, N-glycosylated protein loop

To investigate the role of the IR co-receptor LBD, we first examined the sequence of this region for any unusual structural features. As in iGluRs, the LBD of IRs consists of a ‘Venus fly trap’-like structure formed by two separate lobes (S1 and S2), which are separated in the primary sequence by the ion channel domain (Figure 1A). We generated a multiprotein sequence alignment of the predicted LBD of diverse *D. melanogaster* IRs, including the co-receptors IR8a and IR25a, various tuning IRs, as well as selected iGluRs. This alignment revealed the presence of a stretch of ~30-35 amino acids near the beginning of the S2 domain in IR8a and IR25a that are not aligned to either tuning IR or iGluR sequences (Figure S1). We termed this region the ‘co-receptor extra loop’ (CREL) (Figure 1A). The CREL is highly conserved across orthologous co-receptors from diverse species (Figure 1B and Figure S2) and, consistent with the overall relatedness of the co-receptors, the IR8a and IR25a CRELs share several characteristics, including the presence of short alpha-helical and beta-sheet regions and a single consensus N-glycosylation target motif (NXS/T) (Figure 1B and Figure S2). Notably, although the N-glycosylation site motif is conserved in all CREL sequences in IR8a and IR25a, the precise position varies by precisely four amino acids in a small subset of orthologues (Figure 1B and Figure S2). As the motif is located in an alpha-helical region, this displacement would be predicted to maintain the site on the same face of this helix.

We next sought to determine whether the CREL is N-glycosylated. Because of the limited quantities of protein we could obtain from tissues *in vivo*, we used HEK293 cells to express and purify recombinant IR8a LBD (corresponding to the termite *Zootermopsis nevadensis* sequence). We split the sample in two, treated one with peptide-N-glycosidase F (PNGase F), and subjected both to SDS-PAGE with in-gel tryptic digestion before analysing them using liquid chromatography-tandem mass spectrometry (LC MS/MS). When treated with PNGase F, N-linked glycans are removed from glycosylated asparagine residues, which become deamidated to aspartic acid (resulting in an increase in peptide mass by 1 mass unit) [30]. Thus, an increase in the abundance of tryptic peptides containing aspartic acid following PNGase F treatment is indicative that these sequences originally contained a glycosylated asparagine. By contrast, such treatment does not affect peptides containing unmodified asparagines, whose abundance should therefore remain unchanged. For the IR8a LBD, we identified a tryptic peptide (m/z 648.2984^2+^) whose peak intensity increased ~1000-fold after PNGase F treatment (Figure S3); subsequent analysis unambiguously determined its sequence as DITLN*SSSDQSK (where N* refers to a deamidated asparagine residue), which corresponds to the predicted N-glycosylation site of the CREL (Figure S3A and Figure 1B). By contrast, an adjacent tryptic peptide (m/z 676.3276^2+^; sequence N*AEDVLYNVWK) had similar abundance in the untreated and PNGase F-treated samples (Figure S3B), indicating that this sequence is not N-glycosylated. These data indicate that the predicted CREL N-glycosylation site does indeed bear a sugar modification.

### The CREL glycosylation site has a selective role in subcellular trafficking

To determine whether the CREL and the CREL N-glycosylation site are required for IR function *in vivo*, we focused on *D. melanogaster* IR8a, because the tuning receptor partners of this co-receptor (i.e., acid-sensing IRs in the antenna, the main olfactory organ of insects) are better understood than for IR25a [10, 11, 27, 31]. We generated transgenes encoding EGFP-tagged mutant versions of IR8a bearing either a small deletion of the CREL (removing T658-D681) or a point mutant that disrupts the N-glycosylation (N669Q), as well as a wild-type IR8a control. These transgenes *(UAS-EGFP:Ir8a^wt^*, *UAS-EGFP:Ir8a*^Δ*CREL*^ and *UAS-EGFP:Ir8a^N669Q^)* were inserted at the same location in the *D. melanogaster* genome to eliminate any positional effects on their expression.

We first expressed these transgenes in olfactory sensory neurons (OSNs) under the control of the *Ir8a-Gal4* driver [27]. OSN dendrites are housed within cuticular sensilla that cover the external surface of the antenna as well lining the sacculus, an internal multichambered pocket (Figure 1C-D). We focused our attention initially on the subpopulation of IR8a-positive sacculus OSNs that co-express the tuning receptor IR64a [12], because the larger soma and dendrites of these neurons – compared to other IR8a-expressing OSNs that innervate coeloconic sensilla [1] – facilitates observation of the subcellular distribution of the EGFP fusion proteins.

EGFP:IR8a^wt^ displayed an identical distribution to endogenous IR8a and IR64a in the soma, inner dendritic segment and sensory cilia of the sacculus neurons (Figure 1E) [27]. By contrast, EGFP:IR8a^ΔCREL^ was detected only in the soma and, rarely, the inner segment, but not in the sensory cilia where endogenous IR8a and IR64a are found (Figure 1E). This result indicates a critical role for the CREL in protein folding, complex assembly and/or subcellular localisation. EGFP:IR8a^N669Q^ displayed a similar localisation pattern to EGFP:IR8a^wt^ (Figure 1E).

As these neurons also express endogenous IR8a, we next expressed these transgenes in an *Ir8a* mutant background. As observed previously, EGFP:IR8a^wt^ localised normally while EGFP:IR8a^ΔCREL^ failed to localise to the sensory cilia, and appeared to be destabilised (Figure 1F). Unexpectedly, in the absence of wild-type IR8a, EGFP:IR8a^N669Q^ also did not localise to cilia (Figure 1F). These observations indicate that the presence of wild-type endogenous IR8a can complement the localisation defect of EGFP:IR8a^N669Q^. This result has two important implications: first, that within cilia-localised IR complexes, there must be at least two IR8a subunits, providing *in vivo* evidence for the stoichiometry of IR complexes suggested by *in vitro* experiments [27]. Second, the ability of EGFP:IR8a^N669Q^ to localise to sensory cilia in the presence of IR8a indicates that this mutant protein is defective neither in folding nor in its ability to assemble into transport-competent IR complexes. Rather, the post-translational modification site in the CREL must have a selective effect on subcellular trafficking.

### Heterogeneous requirement for the CREL glycosylation site with tuning IRs

To examine the role of the IR8a CREL in trafficking of other tuning IRs, we surveyed the localisation of these three EGFP:IR8a fusion proteins in Ir8a neurons across the antenna. While EGFP:IR8a^ΔCREL^ was never detected in sensory cilia, EGFP:IR8a^N669Q^ could be observed in the endings of some neurons innervating coeloconic sensilla, even in the absence of endogenous IR8a (Figure S4A-B). We considered whether this differential trafficking ability of IR8a^N669Q^ might be related to the tuning IR with which it is co-expressed.

To test this possibility, we used an *in vivo* heterologous expression system, Or22a neurons, which are housed in basiconic sensilla. These do not express endogenous IR8a (or tuning IRs) and because of their larger size compared to coeloconic sensilla OSNs, are more amenable to visualisation of subcellular protein localisation. Neither EGFP:IR8a^wt^ nor EGFP:IR8a^N669Q^ localise to sensory cilia when expressed alone in Or22a neurons ([27] and data not shown), reflecting the dependence of IR8a upon a tuning IR to form transport-competent complexes. We first expressed either of these versions of IR8a together with IR75a. In combination with EGFP:IR8a^wt^, IR75a can localise to the ciliated endings of these neurons (Figure 2A). By contrast, IR75a+EGFP:IR8a^N669Q^ is not detected beyond the inner segment (Figure 2A). This is similar to the failure of endogenously-expressed IR64a to localise with EGFP:IR8a^N669Q^ in sacculus neurons (Figure 1F). We next tested two other IR8a-depending tuning IRs, IR84a and IR75c. Both localised to sensory cilia together with EGFP:IR8a^wt^ but, in contrast to IR64a or IR75a, both also localised with EGFP:IR8a^N669Q^ (Figure 2B-C). Thus the IR8a^N669Q^ mutant reveals an unexpected heterogeneity in the cilia targeting properties of IR complexes, with some (i.e., those containing IR64a or IR75a) critically dependent on the CREL glycosylation site, and others (i.e., those containing IR75c or IR84a) independent of this post-translational modification.

**Figure 2.**
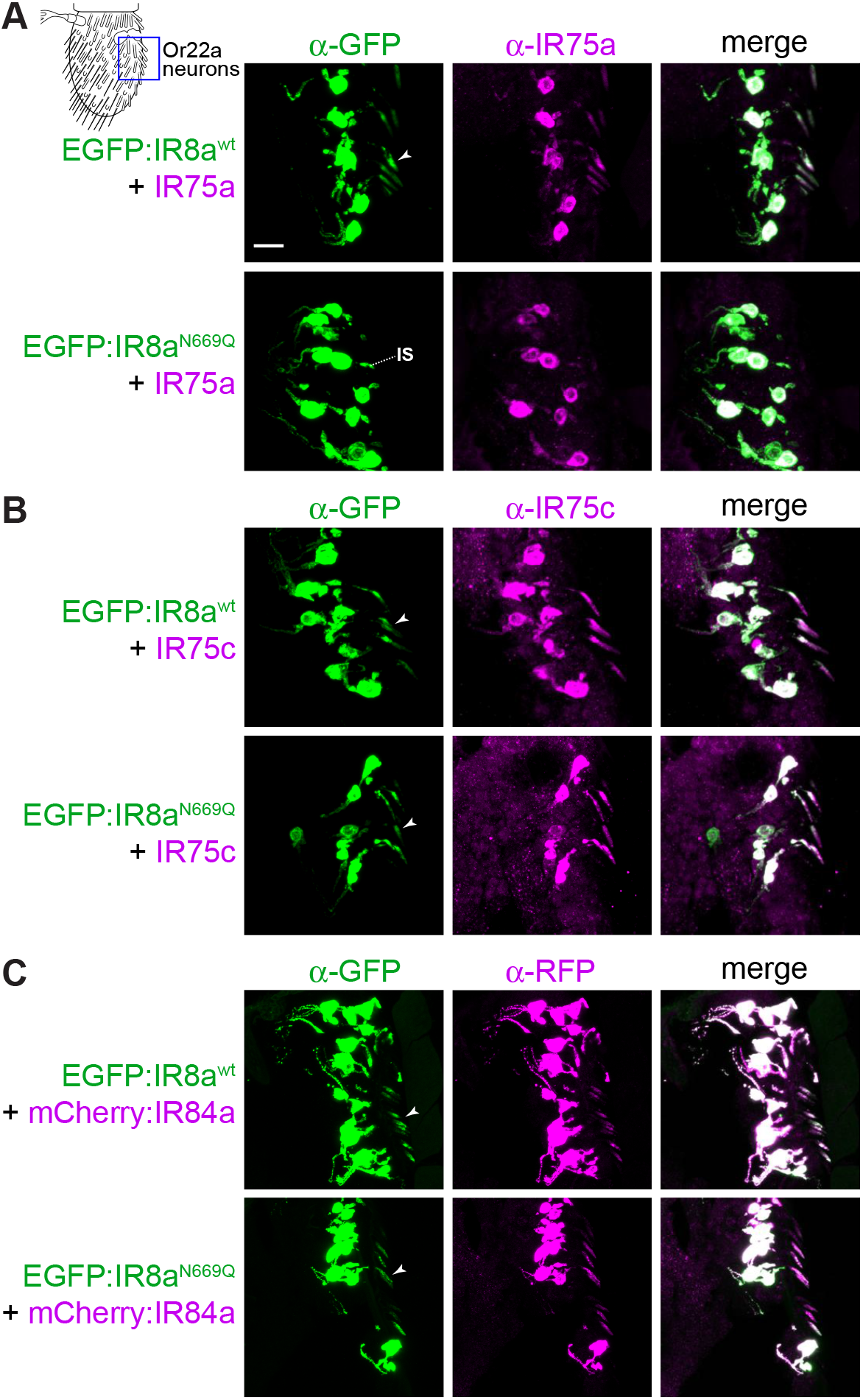
Heterogeneous requirement for IR8a CREL N-glycosylation with tuning IRs. (A)Immunofluorescence with antibodies against EGFP (green) and IR75a (magenta) on antennal sections (representing the field-of-view indicated in the cartoon) of animals expressing the indicated transgenes in Or22a neurons. The arrowheads (in this and other panels) indicate the cilia of Or22a neurons; in neurons expressing EGFP:IR8a^N669Q^ + IR75a (second row), the receptors are not detected in this sensory compartment, remaining restricted to the inner segment (IS). Receptor localisation was determined by overlaying the fluorescence signal onto a bright-field channel (we do not show this channel here because the dense array of cuticular hairs on the antennal surface makes visualisation difficult in a projected image). Note that not all soma have a corresponding ciliated ending in these images, because this is a thin tissue section. Genotypes are of the form: *UAS-EGFP:Ir8a*^*x*^/*UAS-Ir75a;Or22a-Gal4*/+. Scale bar (for all panels in this figure): 10 μm. (B) Immunofluorescence with antibodies against EGFP (green) and IR75c (magenta) on antennal sections of animals expressing the indicated transgenes in Or22a neurons. Genotypes are of the form: *UAS-EGFP:Ir8a*^*x*^/*UAS-Ir75c;Or22a-Gal4/+*. (C) Immunofluorescence with antibodies against EGFP (green) and RFP (magenta) on antennal sections of animals expressing the indicated transgenes in Or22a neurons. Genotypes are of the form: *UAS-EGFP:Ir8a*^*x*^/+;*Or22a-Gal4/UAS-mCherry:Ir84a.*

Our observation that wildtype IR8a can promote cilia transport of IR8a^N669Q^ (Figure 1E-F) raised the question of whether a tuning IR that is targeted to cilia with IR8a^N669Q^ can facilitate the localisation of a tuning IR that cannot, if they are incorporated into a common complex. We tested this possibility in two ways: first, we examined the distribution of IR64a in its own neurons expressing EGFP:IR8a^N669Q^ (but not endogenous IR8a) together with a control receptor (IR75a, which cannot localise to cilia with IR8a^N669Q^ (Figure 2A)) or test receptors (IR75c or IR84a, which can localise with IR8a^N669Q^ (Figure 2B-C)). While coexpression of IR75c or IR84a (but not IR75a) promoted cilia targeting of EGFP: IR8a^N669Q^ (Figure S5A), in no case did this lead to localisation of IR64a to the sensory compartment (Figure S5A). Levels of IR64a were instead substantially reduced upon co-expression of an additional tuning IR. This might be because these ectopically-expressed IRs preferentially combine with EGFP:IR8a^N669Q^, thereby excluding IR64a from associating with IR8a and so destabilising it (as observed previously [27, 31]). Second, in Or22a neurons, we misexpressed IR75a together with EGFP:IR8a^wt^ or EGFP:IR8a^N669Q^, in the absence or presence of IR75c. Unexpectedly, we observed that addition of IR75c led to lower expression and abolished cilia localisation of IR75a co-expressed with EGFP:IR8a^wt^ (Figure S5B), suggesting that IR75c outcompetes – rather than collaborates – with IR75a to form stable, transport-competent complexes. Together these results indicate that different tuning IRs do not readily assemble in a common complex with IR8a.

Finally, we asked what molecular features might explain why some IRs can localise with EGFP:IR8a^N669Q^. IR75c and IR84a are not distinguished from IR64a and IR75a by phylogenetic relatedness [5] or any obvious sequence motifs (data not shown). Given that it is lack of an N-glycosylation site on IR8a that exposed distinct properties of these tuning receptors, we hypothesised that these receptors might have complementary glycosylation sites. The LBD of IR84a contains five putative N-glycosylation motifs: of these, three (N222, N272, N289) are located on the predicted external surface of the domain (based upon comparison of their location with structures of the iGluR LBD (e.g., [32])). We generated mutant versions of IR84a in which all of these sites were mutated (mCherry:IR84a^N222Q,N272Q,N289Q^) and expressed this receptor (or a wild-type mCherry:IR84a control) together with EGFP:IR8a^wt^ or EGFP:IR8a^N669Q^ in Or22a neurons. Cilia localisation was observed in all cases (Figure S6), albeit at varying level, suggesting that IR84a LBD glycosylation is not an essential compensating factor that permits localisation with IR8a^N669Q^. This result also highlights the specific contribution of the IR8a CREL N-glycosylation site to membrane trafficking of IR complexes.

### CREL and CREL N-glycosylation are important for ER export

To determine where in the endomembrane system the trafficking of IR8a^ΔCREL^ and IR8a^N669Q^ is blocked, we visualised the distribution of these EGFP-tagged receptors relative to markers for different organelles: endoplasmic reticulum (ER) (labelled with tdTomato:Sec61β [33]), Golgi apparatus (labelled with γCOP:mRFP [34]) and the cilia transition zone (labelled with antibodies against B9d1 [35]; Figure 1D). We used genetically-encoded markers for the ER and Golgi in order to express them only in the OSNs of interest, thereby avoiding confounding signal from the organelle-rich epidermal cells in the antenna [36].

We first analysed the distribution of EGFP:IR8a variants – co-expressed with IR75a – in Or22a neurons (Figure 3). In these cells, the ER marker displays a prominent perinuclear signal, but also extended up to the base of the transition zone (Figure 3A), suggesting that this organelle is broadly distributed in OSNs. EGFP:IR8a^wt^ displays a similar, though not identical, distribution in the soma and inner dendrite, in addition to its terminal localisation in the sensory cilia (where intracellular organelles are not observed). Both EGFP:IR8a^ΔCREL^ and EGFP:IR8a^N669Q^, though absent from the cilia, display a similar overlap with the ER (Figure 3A). The Golgi marker was found in a few large puncta present primarily in the OSN soma (Figure 3B). These puncta are almost entirely devoid of EGFP:IR8a^wt^ (Figure 3B), suggesting that IRs transit rapidly through this organelle. Alternatively, given the broader distribution of the ER (Figure 3A) the receptors might follow a Golgi-independent route from the ER to the sensory compartment, as described for other cilia membrane proteins [37]. Neither EGFP:IR8a^ΔCREL^ nor EGFP:IR8a^N669Q^ display Golgi localisation, indicating that their inability to localise to sensory cilia is not due to transport arrest in the Golgi. Similarly, we did not detect overlap of any EGFP:IR8a variant and the transition zone marker (Figure 3C), indicating that the wild-type protein passages quickly from inner to outer dendrite, and that the mutant proteins do not become blocked at this stage in their transport to cilia. We repeated these analyses in IR64a-expressing sacculus neurons, and made very similar observations (Figure S7): EGFP:IR8a^ΔCREL^ and EGFP:IR8a^N669Q^ display overlap with the ER marker, but not Golgi or transition zone markers. Together these observations are consistent with a model in which the IR8a^ΔCREL^ and IR8a^N669Q^ fail to localise to sensory cilia because they are trapped in the ER, rather than in later compartments of the transport pathway.

**Figure 3.**
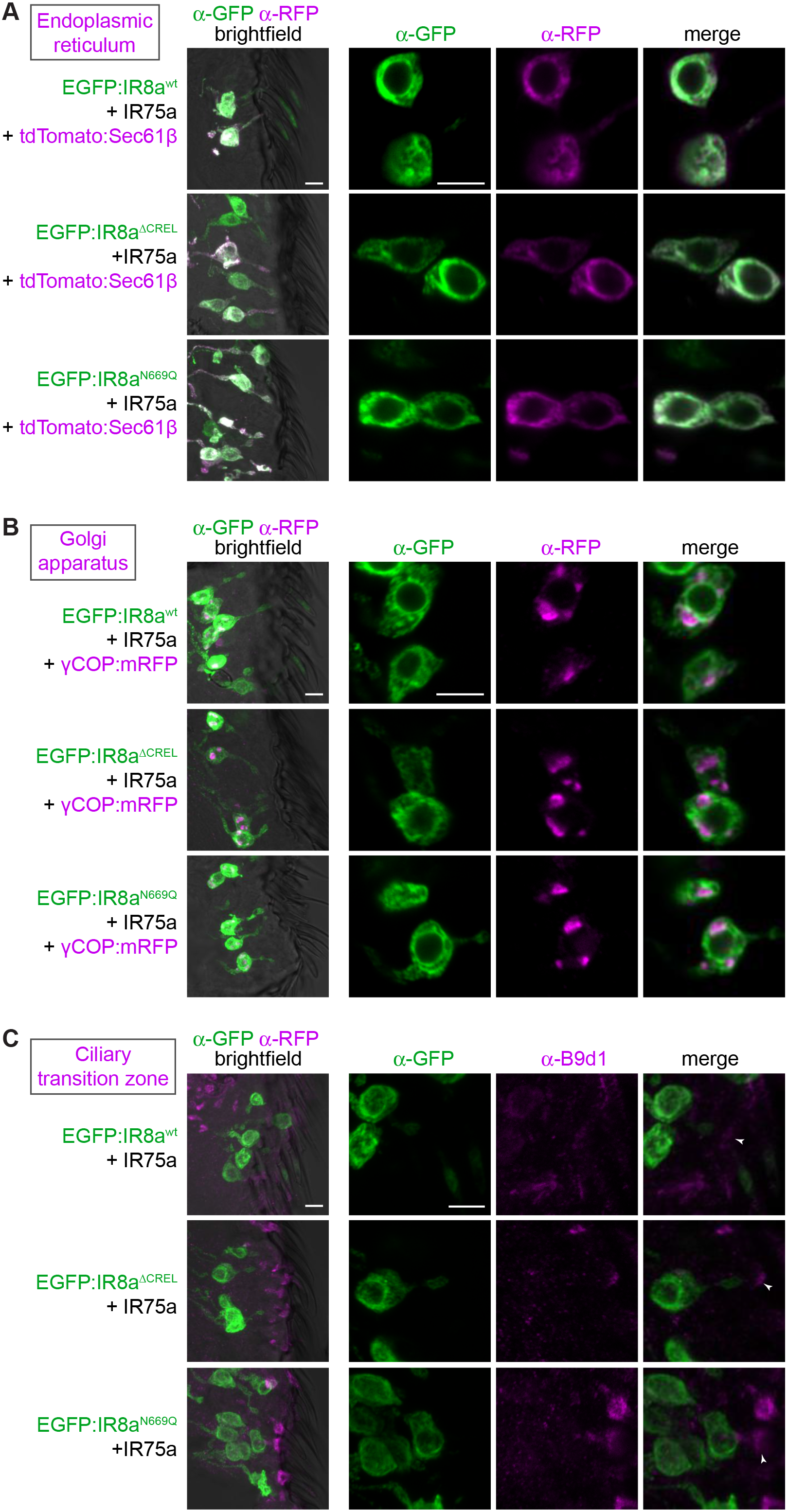
The CREL and CREL glycosylation are important for ER export. (A) Immunofluorescence with antibodies against EGFP (green) and RFP/Tomato (magenta) on antennal sections of animals expressing the indicated transgenes in Or22a neurons. The images on the right are high-magnification, single optical slices taken within the region shown in the lower-magnification view on the left. Genotypes are of the form: *UAS-EGFP:Ir8a*^*x*^/*UAS-Ir75a;Or22a-Gal4/UAS-tdTomato:Sec61β.* Scale bars: 5 μm. (B) Immunofluorescence with antibodies against EGFP (green) and RFP (magenta) on antennal sections of animals expressing the indicated transgenes in Or22a neurons. The images on the right are high-magnification, single optical slices taken within the region shown in the lower-magnification view on the left. Genotypes are of the form: *UAS-EGFP:Ir8a*^*x*^/*UAS-Ir75a;Or22a-Gal4/UAS-γCOP.mRFP.* Scale bars: 5 μm. (C) Immunofluorescence with antibodies against EGFP (green) and B9d1 (magenta) on antennal sections of animals expressing the indicated transgenes in Or22a neurons. The images on the right are high-magnification, single optical slices taken within the region shown in the lower-magnification view on the left (the transition zone is more clearly visible in the lower-magnification images). Genotypes are of the form: *UAS-EGFP:Ir8a*^*x*^/*UAS-Ir75a;Or22a-Gal4*/+. Scale bars: 5 μm.

### CREL N-glycosylation is dispensable for the function of IR complexes

The localisation of a subset of IRs to sensory cilia with IR8a^N669Q^ further allowed us to ask whether this N-glycosylation site also contributes to odour-evoked signalling. Using single sensilla electrophysiological recordings, we measured the responses of Or22a neurons expressing the EGFP-tagged IR8a variants with IR84a to its best-known agonist, phenylacetic acid [38]. IR84a+EGFP:IR8a^wt^ confer robust responses to this odour compared to control neurons that do not express these receptors (Figure 4A-B). Consistent with the lack of cilia localisation, IR84a+EGFP:IR8a^ΔCREL^ do not confer responses above background levels (Figure 4A-B). By contrast, IR84a+EGFP:IR8a^N669Q^ confers phenylacetic acid sensitivity that is indistinguishable from IR84a+EGFP:IR8a^wt^ (Figure 4A-B). We extended this analysis with a second tuning receptor, IR75c, which confers strong responses to propionic acid when co-expressed EGFP:IR8a^wt^, but not EGFP:IR8a^ΔCREL^ (Figure 4C-D). In contrast to the observations with IR84a, IR75c+EGFP:IR8a^N669Q^ co-expression yielded responses that were decreased compared to the IR75c+EGFP:IR8a^wt^ (Figure 4C-D). Together these observations indicate that while the CREL glycosylation is not essential for IR signalling, it may contribute to the function of at least some IR complexes.

**Figure 4.**
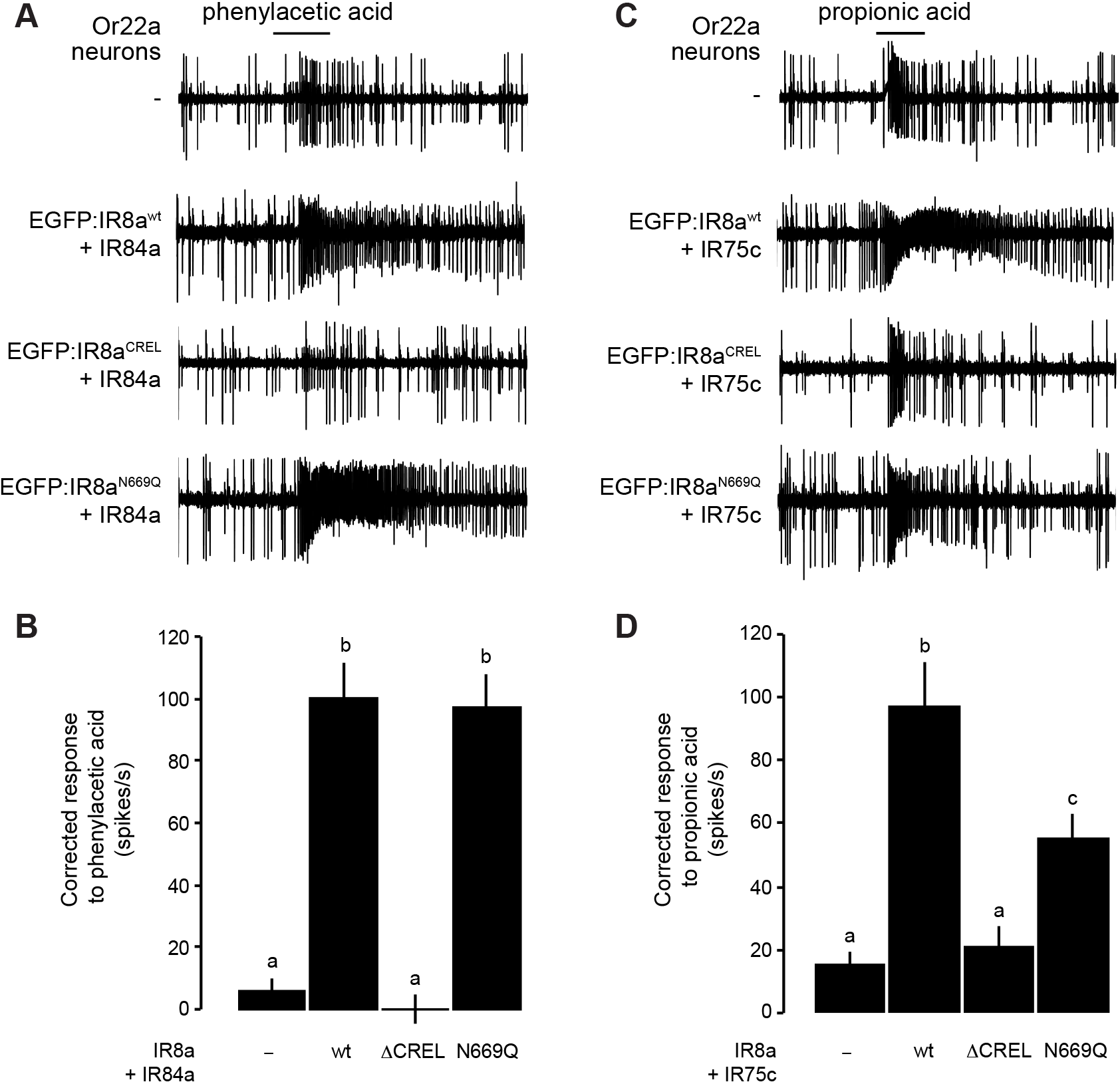
N-glycosylation of the CREL is not essential for odour-evoked IR signalling. (A) Representative traces of responses of Or22a neurons expressing the indicated combinations of IRs, exposed to phenylacetic acid (1% v/v). Genotypes are of the form: *UAS-EGFP:Ir8a*^*x*^/*UAS-Ir84a;Or22a-Gal4*/+, except for the control *(Or22a-Gal4*/+). (B) Quantification of the odour-evoked responses of the genotypes shown in (A). (C) Representative traces of responses of Or22a neurons expressing the indicated combinations of IRs, exposed to propionic acid (1% v/v). Genotypes are of the form: *UAS-EGFP:Ir8a*^*x*^/+;*Or22a-Gal4/UAS-Ir75c,* except for the control *(Or22a-Gal4*/+). (D) Quantification of the odour-evoked responses of the genotypes shown in (C). Mean solvent corrected responses (±SEM; n = 8-12, mixed genders) are shown. Bars labelled with different letters are statistically different (Student t-test with Benjamin and Hochberg correction for multiple comparisons)

### The CREL is likely to be exposed on an external face of an IR heterotetrameric complex

Finally we explored where the CREL is likely to be located within an IR complex. Based upon our *in vivo* evidence (Figure 1E-F), subunit counting analysis *in vitro* [27], and by analogy with the (hetero)tetrameric stoichiometry of iGluRs [2, 3, 29], we reasoned that IR complexes are composed of two IR8a subunits and two tuning subunits. In iGluRs, the extracellular domains of the four subunits form a two-fold axis of symmetry, comprising two with ‘proximal’ ATDs, which contact each other across the axis of symmetry, and two with ‘distal’ ATDs, which do not [39]. The presence of an ATD in IR8a (and IR25a) but not in tuning IRs led us to hypothesise that the IR co-receptor subunits correspond to the iGluR subunits whose ATDs interact. To examine the approximate location of the CREL in such a subunit arrangement, we generated a homology model of a putative IR complex, based upon a structure of the mammalian AMPA receptor GluA2 [32]. In the subunit configuration where IR8a ATDs interact, the CREL is exposed on the external face of the assembled LBDs (Figure 5A). In an alternative model, where IR8a corresponds to iGluR subunits whose ATDs do not contact each other, the CREL is predicted to be buried within the interface between co-receptor and tuning IR subunits (Figure 5B). This latter configuration seems unlikely for two reasons: first, the relatively short (and highly divergent) N-termini of tuning IRs may provide little or no opportunity for specific intersubunit interactions to occur within the upper layer of the complex, a region that is key for assembly in iGluRs [29, 40]. Second, the externally exposed IR8a CREL in the former configuration (Figure 5A) would provide better access to both the N-glycosylation machinery and to (unknown) ER export quality control sensors.

**Figure 5.**
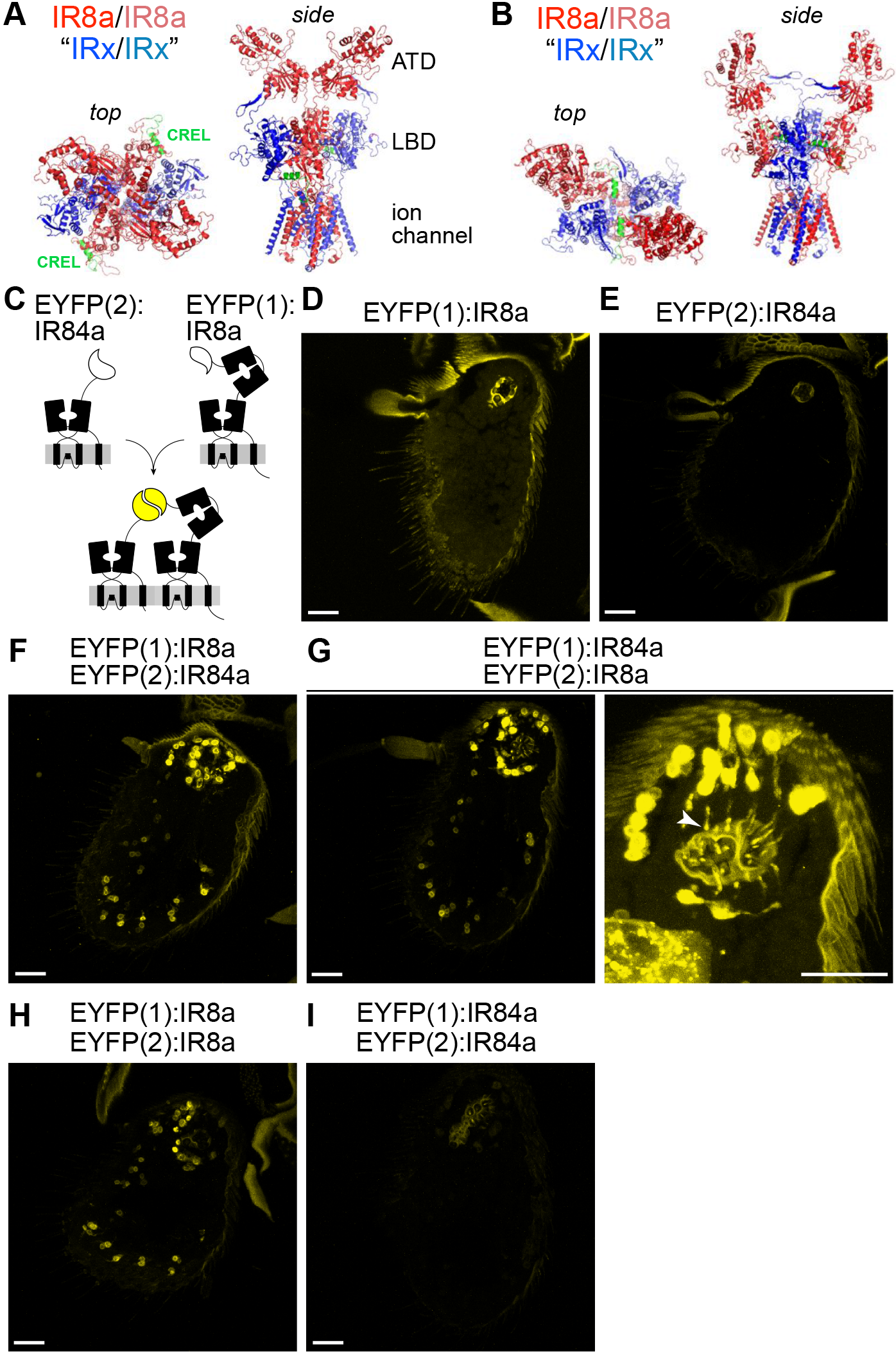
Location of the CREL in a heterotetrameric IR model. (A-B) Two possible configurations of a hypothetical model of an IR heterotetramer of two IR8a subunits (red) and two tuning IR subunits (blue), in which the IR8a ATDs have proximal (contacting) (A) or distal (non-contacting) (B) positions. Top and side views are shown in slightly different orientations to facilitate visibility of the IR8a CREL (green). The structure was built through coarse-grained homology modelling of *D. melanogaster* IR8a on the homotetrameric GluA2 structure [32]; the IR tuning subunits are represented simply by the same IR8a model from which the ATD is deleted. (C) Schematic of the principle of EYFP reconstitution through complex formation and/or close proximity of EYFP fragment:IR fusion proteins. (D-I) Endogenous EYFP fluorescence in antennal sections of animals expressing the indicated combinations of EYFP fragment fusions in Ir8a neurons. The higher magnification sacculus image in (G, right) reveals the cilia localisation of fluorescent signals (arrowhead); here, the gain setting during imaging was increased, resulting in higher cuticular autofluorescence. Genotypes: (D) *UAS-EYFP(1):Ir8a*/+;*Ir8a-Gal4*/+ (E) *UAS-EYFP(2):Ir84a*/+;*Ir8a-Gal4*/+ (F) *UAS-EYFP(1):Ir8a/UAS-EYFP(2):Ir84a;Ir8a-Gal4*/+ (G) *UAS-EYFP(1):Ir84a*/*UAS-EYFP(2):Ir8a;Ir8a-Gal4*/+ (H) *UAS-EYFP(1):Ir8a/UAS-EYFP(2):Ir8a;Ir8a-Gal4*/+ (I) *UAS-EYFP(1):Ir84a/UAS-EYFP(2):Ir84a;Ir8a-Gal4*/+. Scale bars: 20 μm.

To obtain experimental evidence for the configuration of IRs *in vivo*, we used a protein fragment complementation assay with an enhanced yellow fluorescent protein (EYFP) reporter [41, 42]. Complementary and non-associating subfragments of EYFP (EYFP(1) and EYFP(2)) were fused to the N-termini of IR8a and IR84a, respectively, separated by short flexible linkers (Figure 5C), and these proteins were expressed in Ir8a neurons singly or together. Neither fusion protein alone was fluorescent (Figure 5D-E), but upon co-expression we detected a robust EYFP signal in all Ir8a neurons (Figure 5F), indicating direct association or close apposition of IR84a and IR8a. Similar results were obtained when EYFP tags were exchanged on these IRs (Figure 5G). EYFP fluorescence was detected both around the nucleus in the soma and in the ciliated dendritic endings, but not axons, indicating that the complex is likely to form in the endoplasmic reticulum (Figure 5G, right).

In addition to analysis of these heteromeric interactions, we expressed together EYFP(1):IR8a and EYFP(2):IR8a. These fusion proteins also reconstituted EYFP fluorescence (Figure 5H), suggesting the existence of homomeric interactions between two IR8a subunits. We also expressed EYFP(1):IR84a and EYFP(2):IR84a together, but observed that these did not reconstitute a fluorescence signal (Figure 5I). The simplest explanation for this result is that the EYFP fragments on IR84a subunits are not sufficiently close to each other in a tetrameric complex and/or are sterically inhibited from associating due to a ‘barrier’ of the interacting IR8a ATDs. Importantly, the result also provides a negative control that indicates that the EYFP reconstitution observed in the tagged IR84a+IR8a and IR8a+IR8a pairs is likely due to the formation of specific protein complexes, rather than simply their co-expression in the same neuronal membranes. Together these observations are consistent with the model of an IR heterotetramer in which the ATDs of two IR8a subunits are directly apposed (Figure 5A).

### Concluding remarks

Our characterisation of the IR8a CREL has revealed a critical role of the IR coreceptor LBD in regulating intracellular transport from the ER to sensory cilia, a property that is likely to be relevant for the functionally-diverse IR8a-and IR25a-containing sensory receptor complexes. Our data also provide novel insights into the stoichiometry and assembly of IR complexes *in vivo*, supporting a model in which two co-receptor subunits form a ‘core’ – possibly interacting via their ATDs [27] – with which two tuning IR subunits associate in the ER.

The unexpected heterogeneity in the localisation properties of different IR complexes in which N-glycosylation of the CREL is prevented suggests that each tuning/co-receptor complex has a unique conformation for passing a trafficking check-point in the ER: in some cases, the CREL N-glycosylation site is a key part of the signal permitting exit from this organelle, while in others it is dispensable. In the absence of obvious molecular features that can account for the distinction of different IRs, we speculate that the observed heterogeneity is related to the conformational flexibility of the LBDs within an IR complex, as this is a property of iGluR LBDs that influences ER export [43].

Such context-dependent trafficking properties in the IR family are reminiscent of the variable dependence of different mammalian Odorant Receptors on accessory proteins for ER exit [44]. This heterogeneity may reflect the conflict that exists during the evolution of divergent chemosensory receptor families, as individual members are under selective pressure both to maintain conserved cellular properties (i.e., trafficking to sensory cilia) and to acquire novel sensory-detection capacities.

## Acknowledgements

We acknowledge Benoîte Bargeton and Sabine Mentha for preliminary experiments and generation of plasmid constructs. We thank Bénédicte Durand for antibodies, the Bloomington *Drosophila* Stock Center (NIH P40OD018537) for flies, and members of the Benton laboratory for comments on the manuscript. L.L.P.-G. was supported by a FEBS Long-Term Fellowship. This work was supported by the University of Lausanne, ERC Starting Independent Researcher and Consolidator Grants (205202 and 615094) to R.B., an HFSP Young Investigator Award (RGY0073/2011) to R.B. and R.J., the SNSF Nano-Tera Envirobot project (20NA21_143082) to R.B., and the National Institutes of Health (R01GM074830) to L.H.

## Author contributions

L.A. performed all experiments except for Figure S3 (H.P., C.G., L.H. and R.J.) and Figure 4 (L.L.P.-G.). R.B. generated sequence alignments and protein models, and generated some plasmids constructs. All authors contributed to experimental design, analysis and interpretation of results. R.B. directed the project and wrote the paper with input from other authors.

## Materials and Methods

### Bioinformatics

Alignments were made with MUSCLE [45] and visualised in Jalview 2.9.0b2 [46]. Secondary structure predictions were made using Quick2D [47]. The IR8a homology model was built using SWISS-MODEL [48], with the *R. norvegicus* GluA2 structure (PDB: 6DLZ [32]) as template.

### Molecular biology

Deletions and point mutations in *Ir8a* and *Ir84a* coding sequences (lacking the region encoding the endogenous signal sequence, as described previously [27]) were introduced by standard PCR-based deletion and site-directed mutagenesis methods. Wild-type and mutant *Ir8a* sequences were subcloned into *pUAST-EGFP* attB, which contains the calreticulin signal sequence (SS) fused to EGFP [27]. Wild-type and mutant *Ir84a* sequences were subcloned into an equivalent *pUAST-mCherry* attB [27]. Similarly, transgenes for EYFP protein fragment complementation were generated by joining sequences encoding SS:EYFP(1) or SS:EYFP(2) [42] to *Ir8a* or *Ir84a* sequences with an intervening short linker (encoding [GGGGS]_2_) in *pUAST attB* [5].

### Biochemistry

*Protein purification:* the sequence of *ZnevIr8a^S1S2^* was synthesised by GENEWIZ, and encodes the S1 domain (residues A366-P490) and the S2 domain (residues P608-N781) connected by a GTGT peptide. This sequence was cloned into a pcDNA vector for mammalian cell expression with a human IL2 signal sequence (MYRMQLLSCIALSLALVTNS), a 9xHis-tag, and a Factor Xa cleavage site added to its N-terminus. The protein was expressed and secreted from FreeStyle HEK 293-F cells (ThermoFisher) and purified directly from cell culture medium using Ni2+−NTA resins. The protein was eluted from the resins with high concentration of imidazole and further purified by a Superdex-200 size-exclusion column in a buffer containing 50 mM Tris, pH 8.0, and 400 mM NaCl.

*Peptide sequencing by LC MS/MS:* tryptic peptides were analysed by LC MS/MS using an LTQ-Orbitrap XL MS (Thermo Fisher, San Jose, CA) coupled on-line with an Easy-nLC 1000 (Thermo Fisher, San Jose, CA) [49]. Each MS/MS experiment consisted of one MS scan in FT mode (350-1800 m/z, resolution of 60,000 at m/z 400) followed by ten data-dependent MS/MS scans in IT mode with normalised collision energy at 29%. Protein identification and characterisation were performed by database searching using the Batch-Tag within the developmental version of Protein Prospector (v 5.17.0) Protein Prospector [49] against a targeted database consisting of *Znev*IR8a sequences. The mass accuracies for parent ions and fragment ions were set as ± 20 ppm and 0.6 Da, respectively. Trypsin was set as the enzyme, and a maximum of two missed cleavages were allowed. Protein N-terminal acetylation, methionine oxidation, N-terminal conversion of glutamine to pyroglutamic acid, and asparagine deamidation were set as variable modifications. Peptide relative abundances were evaluated based on extracted chromatograms of the selected ions during MS scans.

### *Drosophila* strains

Flies were maintained at 25°C in 12 h light: 12 h dark conditions. We used the following published *D. melanogaster* strains: *Ir8a*^*1*^, *Ir8a-Gal4* [27], *Or22a-Gal4* [50], *UAS-Ir75a* [10], *UAS-Ir75c* [11], *UAS-Ir84a* [1], *UAS-tdTomato:Sec61β* (BSDC#64747) [33], *UAS-γCOP:mRFP* (BDSC#29714) [34]. New transgenic flies were generated by Genetic Services Inc. or BestGene Inc., via the phiC31 site-specific integration system, using the *attP40* and *attP2* landing site strains [51] for insertions on chromosome 2 and 3, respectively. Genotypes are provided in the figure legends.

### Immunohistochemistry and imaging

Primary antibodies: guinea pig anti-IR8a (1:1000) [27], rabbit anti-IR64a (1:1000) [12], rabbit anti-IR75a (1:1000) [10], rabbit anti-IR75c (1:200) [11], guinea pig anti-B9d1 (1:2500) [35], anti-GFP mouse (1:1000) (Invitrogen A11120), anti-RFP (1:1000) (Abcam ab62341). Secondary antibodies: Alexa488 anti-mouse (1:1000) (A11029 INVITROGEN), Cy3 anti-guinea pig (1:1000) (Jackson Immunoresearch 106-166-003), Cy3 anti-rabbit (1:1000) (MILAN Analytica AG 111-165-144 0).

Microscopy was performed using a Zeiss LSM 710 Inverted Laser Scanning Confocal Microscope. Confocal images were processed with Fiji [52]. For all experiments presented, antennae from a minimum of 18 flies from at least three independent immunostainings from two separate genetic cross were examined, except in Figure S5A, where only 6-12 flies from 1-2 separate crosses were examined.

### *In vivo* electrophysiology

Single sensillum extracellular recordings were performed and analysed essentially as described previously [53], using odour cartridges assembled as detailed in [54]. Phenylacetic acid (CAS #103-82-2) and propionic acid (CAS: 7909-4) were from Sigma-Aldrich and were of the highest purity available. Odorants were used at 1% (v/v) in double-distilled H_2_O.

## Supplementary Figure Legends

**Figure S1.**
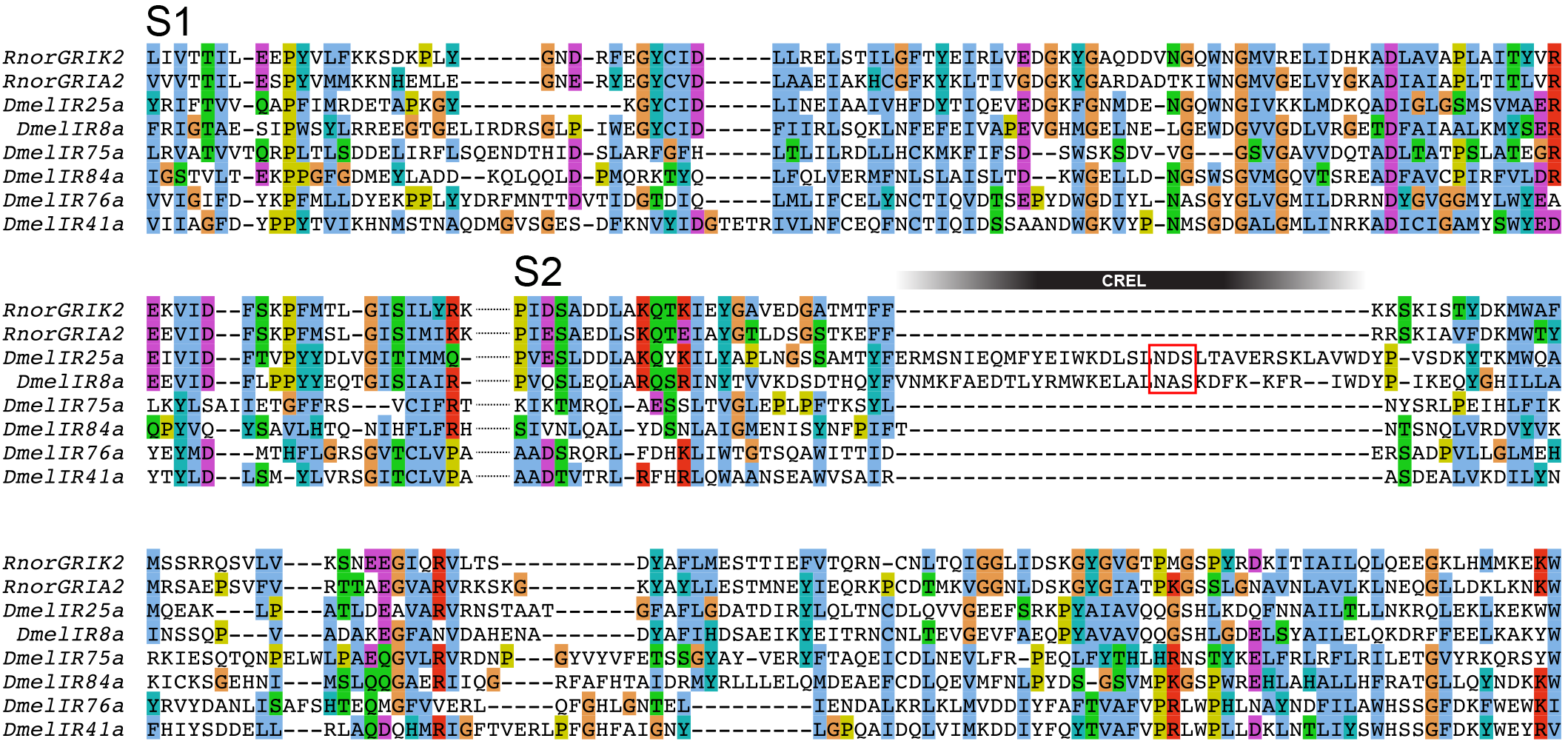
Alignment of IR and iGluR LBDs. Multiple sequence alignment of the predicted ligand-binding domain sequence from the indicated *R. norvegicus* iGluRs and *D. melanogaster* IRs. The approximate position of the CREL is indicated, and the conserved N-glycosylation site within this sequence is highlighted with a red box.

**Figure S2.**
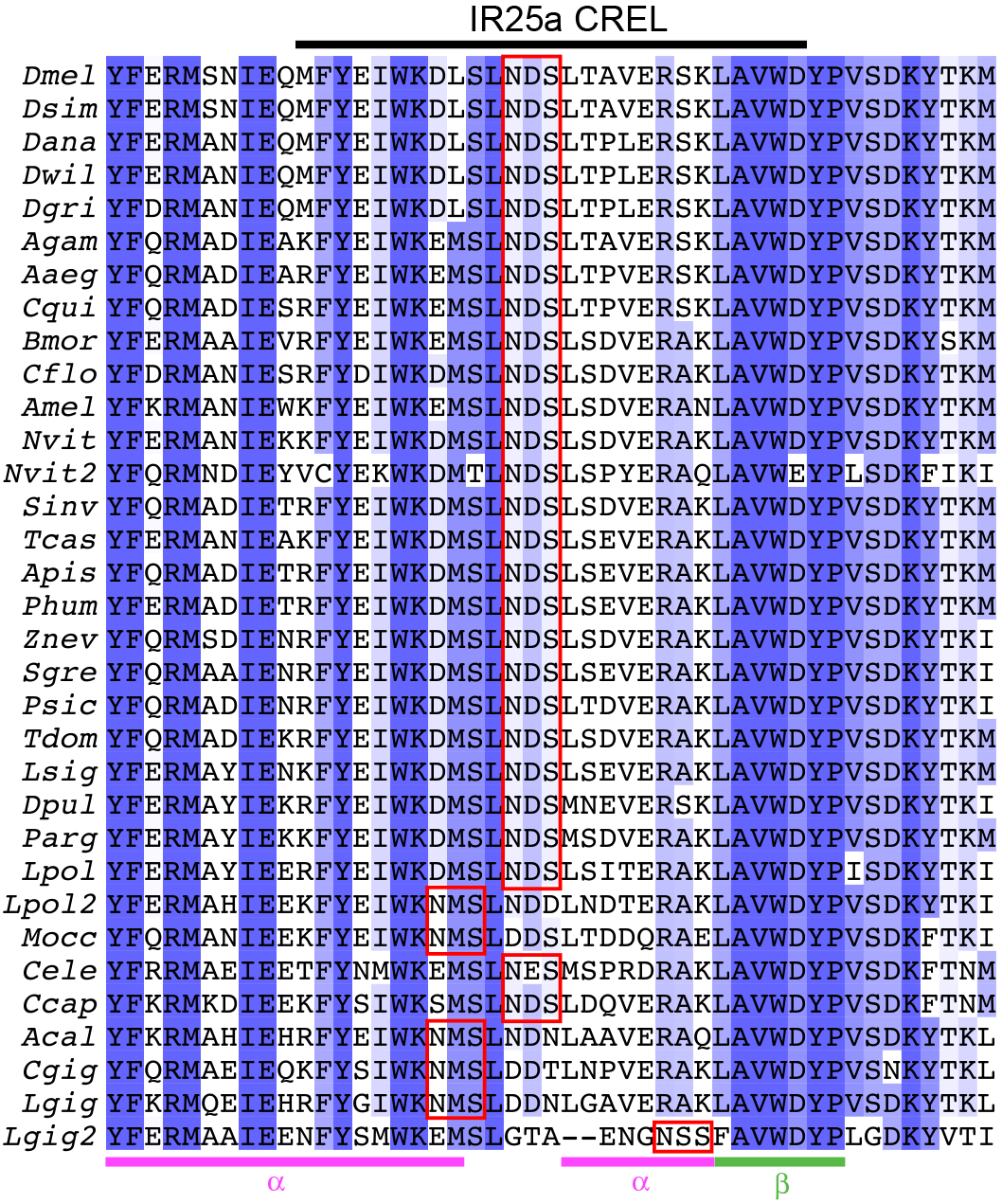
IR25a CREL alignment. Alignment of the protein sequence spanning the CREL in IR25a orthologues from the indicated species. Predicted N-glycosylation sites are highlighted with red boxes and predicted secondary structure is shown below the alignment. Species (top-to-bottom): *Drosophila melanogaster*, *Drosophila simulans*, *Drosophila ananassae*, *Drosophila willistoni*, *Drosophila grimshawi*, *Culex quinquefasciatus*, *Aedes aegpti*, *Anopheles gambiae*, *Bombyx mori*, *Tribolium castaneum*, *Camponotus floridanus*, *Solenopsis invicta*, *Apis mellifera*, *Nasonia vitripennis* (two orthologues), *Zootermopsis nevadensis*, *Acyrthosiphon pisum*, *Pediculus humanus*, *Schistocerca gregaria*, *Daphnia pulex*, *Panulirus argus*, *Limulus polyphemus* (two orthologues), *Metaseiulus occidentalis*, *Caenorhabditis elegans*, Capitella *capitata*, *Aplysia californica*, *Crassostrea gigas*, *Lottia gigantea* (two orthologues).

**Figure S3.**
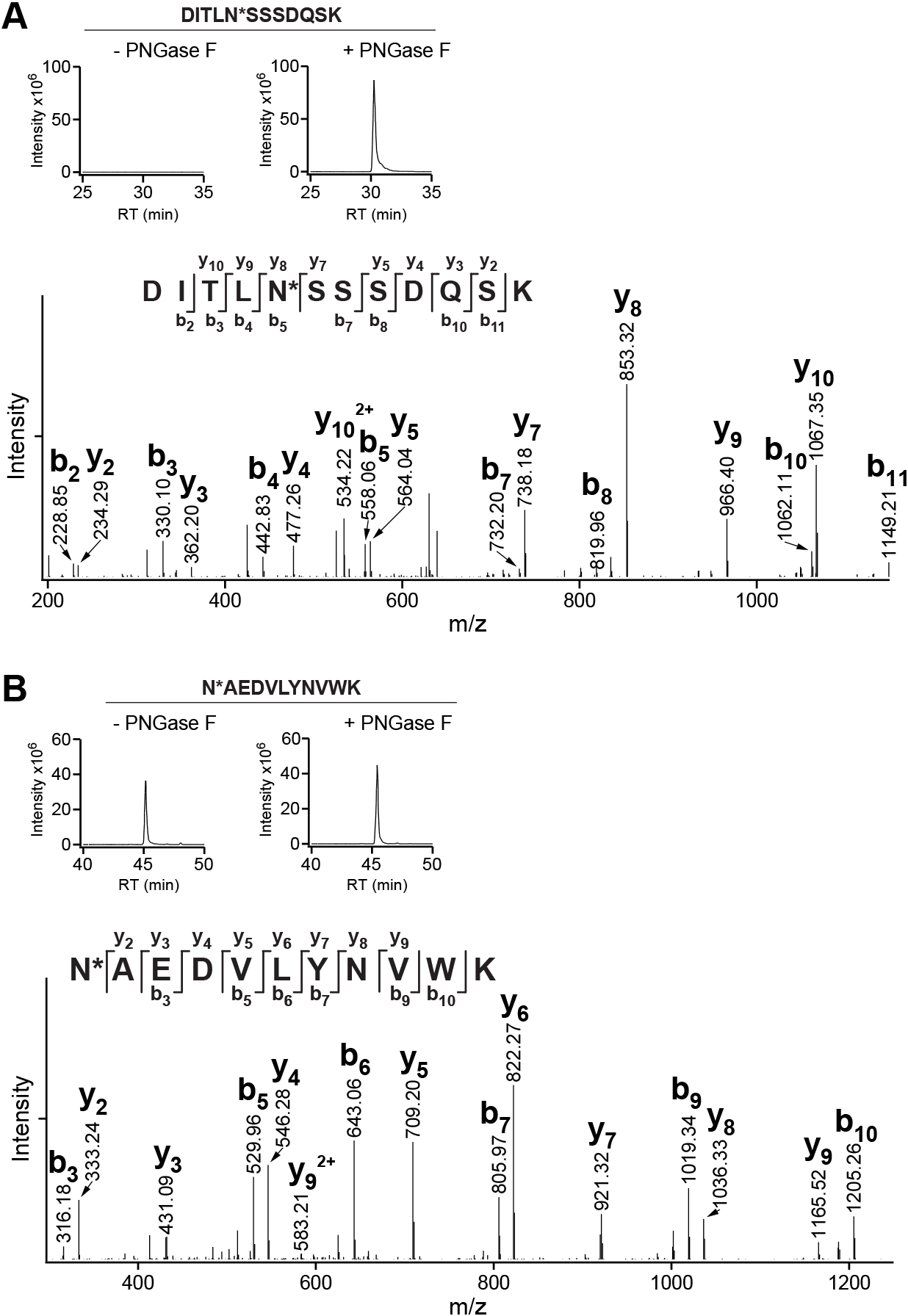
The IR8a CREL contains a single N-linked glycosylation site. (A) Top: extracted ion chromatograms of a *Zootermopsis nevadensis (Znev)* IR8a tryptic peptide containing a deamidated asparagine (N*) (m/z 648.2984^2+^) before and after PNGase F treatment; the abundance of this peptide increases 1000-fold after treatment. Bottom: MS/MS spectrum identifying the corresponding peptide (DITLN*SSSDQSK, within the CREL (Figure 1B)). (B) Top: extracted ion chromatograms of a *Znev*IR8a tryptic peptide containing a deamidated asparagine (m/z 676.3276^2+^) before and after PNGase F treatment Bottom: MS/MS spectrum identifying the corresponding peptide sequence (N*AEDVLYNVWK), which lies at the beginning of the CREL sequence (Figure 1B). In this peptide, the deamidated terminal asparagine does not indicate an N-glycosylated residue, because peptide abundance is similar with and without PNGase F treatment, and mostly likely reflects an artefact of MS sample preparation.

**Figure S4.**
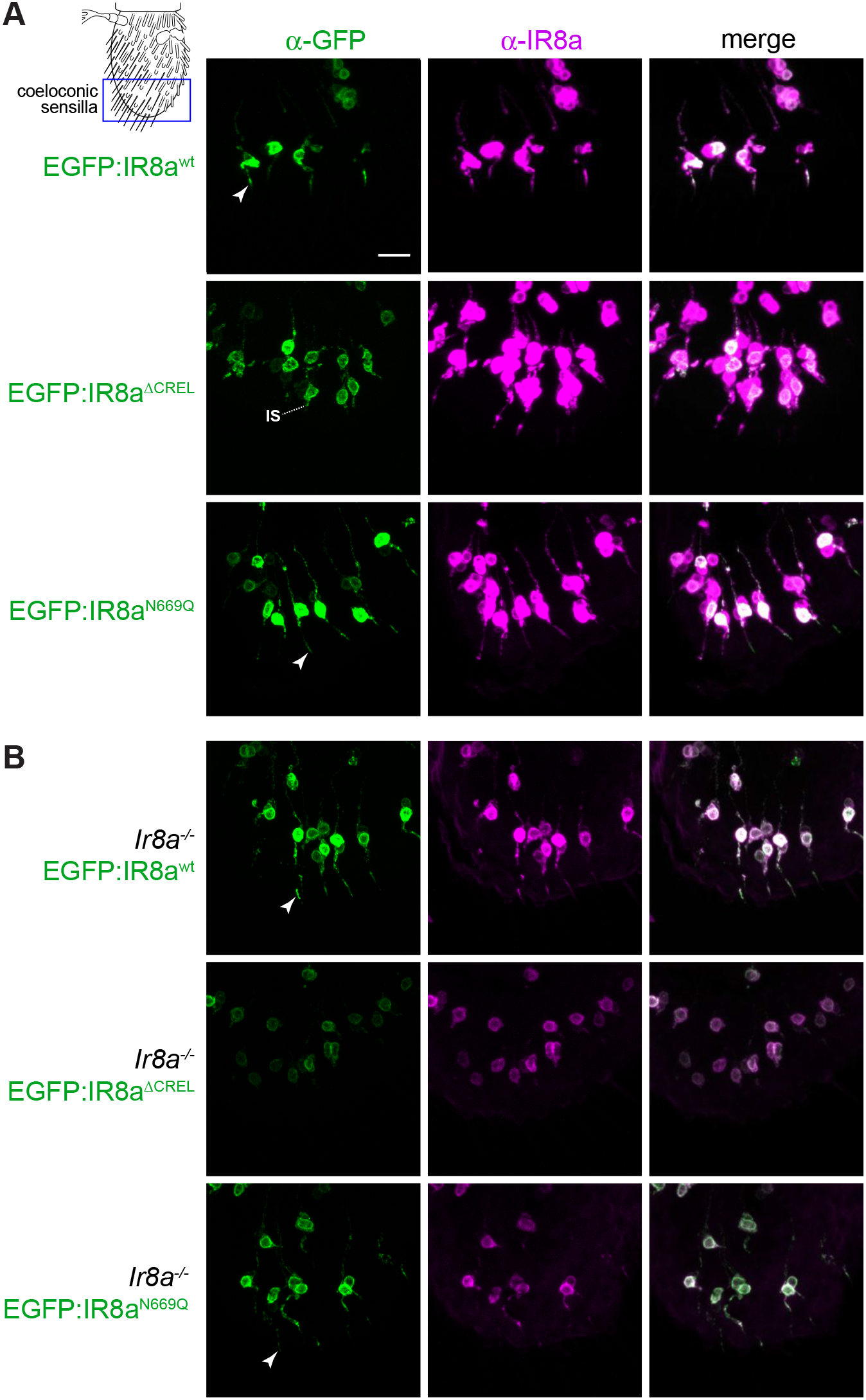
Heterogeneous localisation properties of IR8a^N669Q^ in coeloconic sensilla. (A) Immunofluorescence with antibodies against EGFP (green) and IR8a (magenta) on antennal sections of animals expressing the indicated transgenes in Ir8a neurons. Genotypes are of the form: *Ir8a-Gal4*/*UAS-EGFP:Ir8a*^*x*^. Arrowheads mark examples of sensilla in which receptors are present in the OSN cilia; this was determined by overlaying the fluorescence signal onto a bright-field channel (we do not show this channel here because the dense array of cuticular hairs on the antennal surface makes visualisation difficult in a projected image). EGFP:IR8a^ΔCREL^ does not traffic beyond the inner segment (IS). Scale bar (for all panels in this figure): 10 μm. (B) Immunofluorescence with antibodies against EGFP (green) and IR8a (magenta) on antennal sections of animals expressing the indicated transgenes in Ir8a neurons in an *Ir8a* mutant background. Genotypes are of the form: *Ir8a^1^*/*Y;Ir8a-Gal4/UAS-EGFP:Ir8a*^*x*^. Arrowheads mark examples of sensilla in which receptors are present in the OSN cilia.

**Figure S5.**
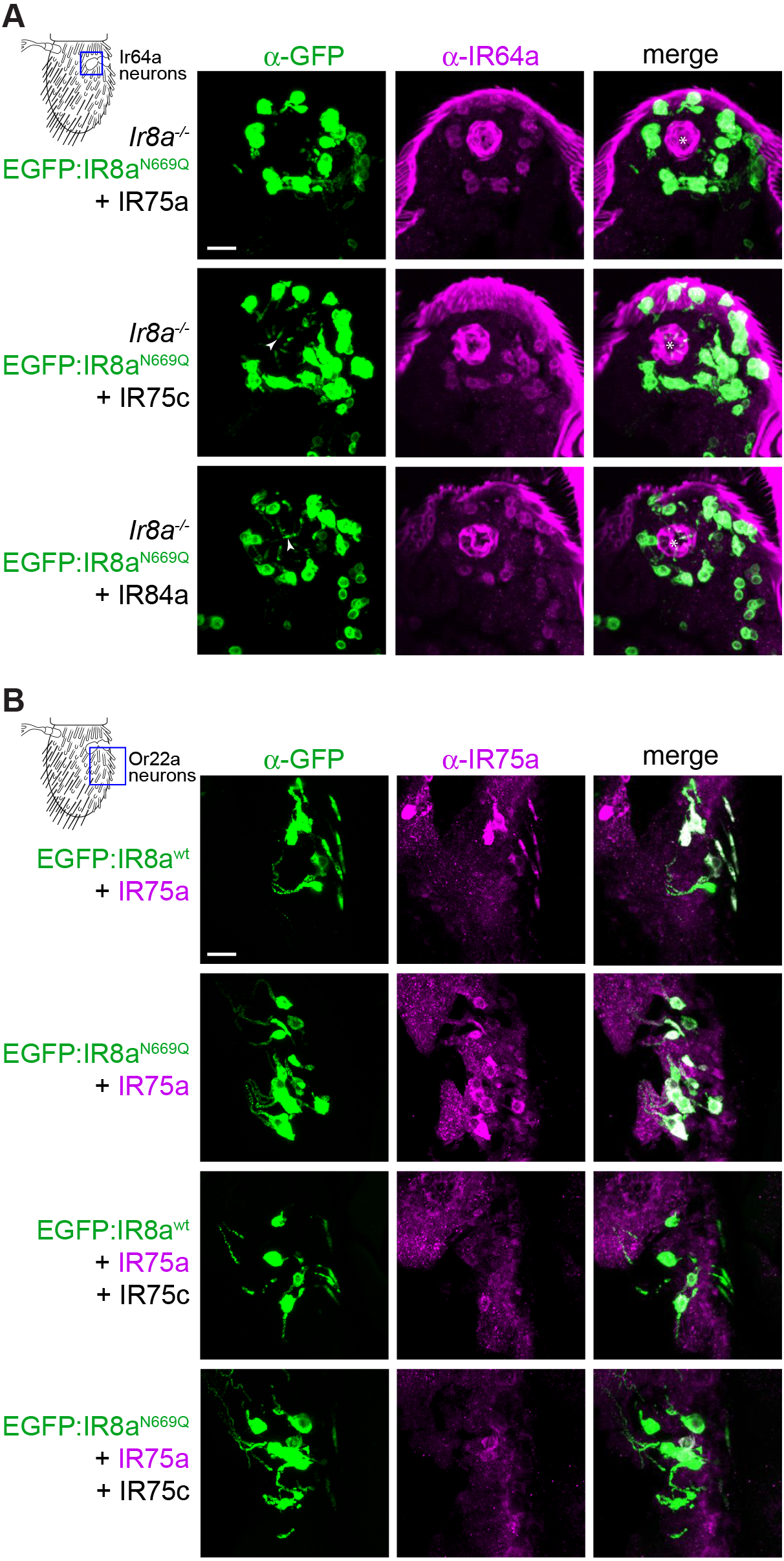
Tuning IRs compete for, rather than combine together with, IR8a^N669Q^. (A) Immunofluorescence with antibodies against EGFP (green) and IR64a (magenta) on antennal sections of animals expressing the indicated transgenes in *Ir8a* neurons in an *Ir8a* mutant background. Genotypes are of the form: *Ir8a^1^/Y;UAS-EGFP:Ir8a^N669Q^/UAS-IrXX;Ir8a-Gal4*/+. The white asterisks in the right-hand panels indicate the central cavity of sacculus chamber 3 into which the OSN ciliated dendrites project. Due to the weak expression of IR64a in these tissues (compared to, for example, Figure 1E-F), the gain setting during imaging was increased, resulting in high cuticular autofluorescence in the magenta channel, which reveals both the antennal surface and the lining of the sacculus (resembling a ‘ball’ due to the sectioning). The arrowheads in the left-hand panels mark the ciliated endings of neurons containing EGFP:IR8a^N669Q^ (but not IR64a). Scale bar (for all panels in this figure): 10 μm. (B) Immunofluorescence with antibodies against EGFP (green) and IR75a (magenta) on antennal sections of animals expressing the indicated transgenes in Or22a neurons. Genotypes are of the form: *UAS-EGFP:Ir8a*^*x*^/+;*Or22a-Gal4/UAS-Ir75a* (top two rows) and *UAS-EGFP:Ir8a*^*x*^/*UAS-Ir75c;Or22a-Gal4/UAS-Ir75a* (bottom two rows).

**Figure S6.**
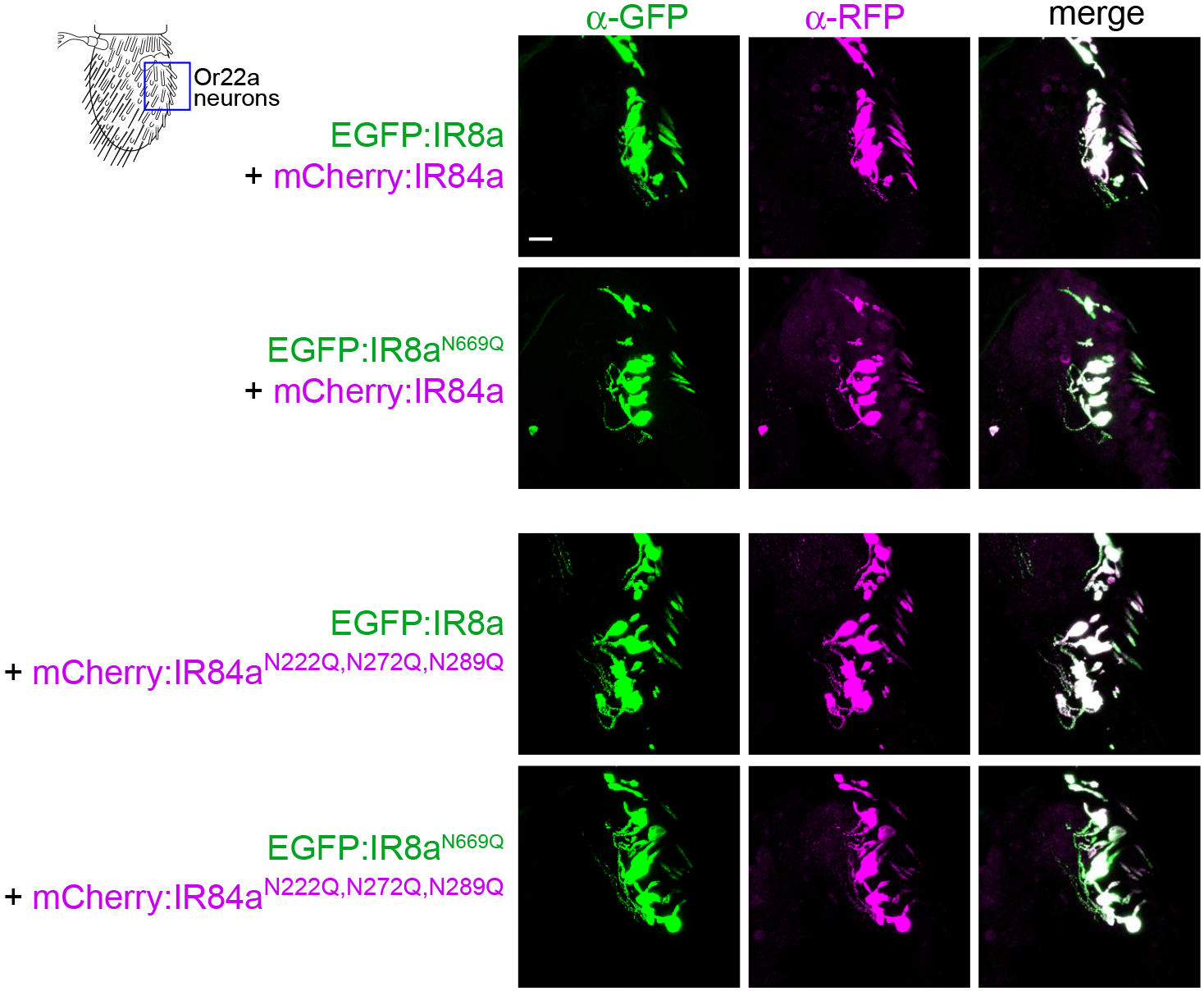
Predicted IR84a N-glycosylation sites do not contribute to cilia localisation of IR complexes in the presence or absence of IR8a CREL glycosylation. Immunofluorescence with antibodies against EGFP (green) and RFP (magenta) on antennal sections of animals expressing the indicated transgenes in Or22a neurons. Genotypes are of the form: *UAS-EGFP:Ir8a*^*x*^/+;*Or22a-Gal4/UAS-mCherry:Ir84a*^*x*^. Scale bar (for all panels in this figure): 10 μm.

**Figure S7.**
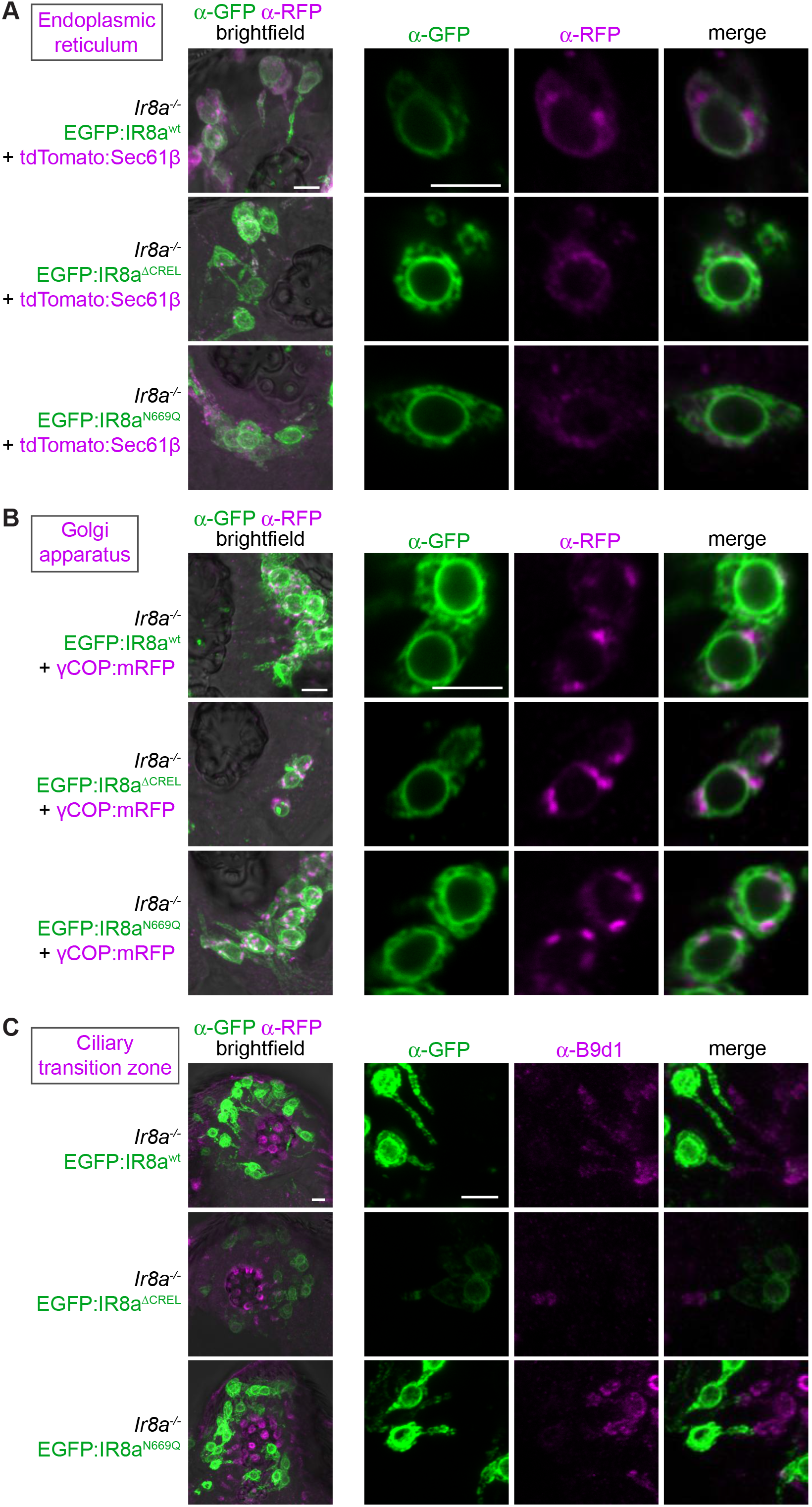
CREL and CREL glycosylation are important for ER export. (A) Immunofluorescence with antibodies against EGFP (green) and RFP/Tomato (magenta) on antennal sections of animals expressing the indicated transgenes in Ir8a neurons. The images on the right are high-magnification, single optical slices taken within the region shown in the lower-magnification view on the left. Genotypes are of the form: *Ir8a*^*1*^/*Y;Ir8a-Gal4/UAS-EGFP:Ir8a*^*x*^;*UAS-tdTomato:Sec61*β/+. Scale bars: 5 μm. (B) Immunofluorescence with antibodies against EGFP (green) and RFP (magenta) on antennal sections of animals expressing the indicated transgenes in Ir8a neurons. The images on the right are high-magnification, single optical slices taken within the region shown in the lower-magnification view on the left. Genotypes are of the form: *Ir8a*^*1*^/*Y;Ir8a-Gal4/UAS-EGFP:Ir8a*^*x*^;*UAS-γCOP:mRFP*/+. Scale bars: 5 μm. (C) Immunofluorescence with antibodies against EGFP (green) and B9d1 (magenta) on antennal sections of animals expressing the indicated transgenes in Ir8a neurons. The images on the right are high-magnification, partial stacks of optical slices taken within the region shown in the lower-magnification view on the left. Genotypes are of the form: *Ir8a*^*1*^/*Y;Ir8a-Gal4/UAS-EGFP:Ir8a*^*x*^. Scale bars: 5 μm.

